# An fMRI study of phrase structure and subject island violations

**DOI:** 10.1101/2024.05.05.592579

**Authors:** William Matchin, Diogo Almeida, Gregory Hickok, Jon Sprouse

**Affiliations:** Dept. of Communication Sciences and Disorders, University of South Carolina; Program in Psychology, New York University Abu Dhabi; Dept. of Cognitive Sciences and Dept. of Language Science, University of California, Irvine

**Keywords:** subject islands, phrase structure violations, acceptability judgments, fMRI, syntax

## Abstract

In principle, functional neuroimaging provides uniquely informative data in addressing linguistic questions, because it can indicate distinct processes that are not apparent from behavioral data alone. This could involve adjudicating the source of unacceptability via the different patterns of elicited brain responses to different ungrammatical sentence types. However, it is difficult to interpret brain activations to syntactic violations. Such responses could reflect processes that have nothing intrinsically related to linguistic representations, such as domain-general executive function abilities. In order to facilitate the potential use of functional neuroimaging methods to identify the source of different syntactic violations, we conducted an fMRI experiment to identify the brain activation maps associated with two distinct syntactic violation types: phrase structure (created by inverting the order of two adjacent words within a sentence) and subject islands (created by extracting a wh-phrase out of an embedded subject). The comparison of these violations to control sentences surprisingly showed no indication of a generalized violation response, with almost completely divergent activation patterns. Phrase structure violations seemingly activated regions previously implicated in verbal working memory and structural complexity in sentence processing, whereas the subject islands appeared to activate regions previously implicated in conceptual-semantic processing, broadly defined. We review our findings in the context of previous research on syntactic and semantic violations using event-related potentials. Although our results suggest potentially distinct underlying mechanisms underlying phrase structure and subject island violations, our results are tentative and suggest important methodological considerations for future research in this area.

## 1. Introduction

Acceptability judgments provide the primary data for (generative) syntactic theories (see Schütze, 1996 for a review). The linking hypothesis adopted by (generative) syntacticians is that acceptability is a consciously-accessible percept driven by (most likely, automatic) error signals that arise during sentence processing (Schütze, 1996; Sprouse, 2020). The consensus is that these error signals can derive from multiple distinct sources, such as syntactic violations, semantic violations, discourse/pragmatic violations, intonation violation, working memory capacity overload, sentence processing complexity, etc (Featherston, 2005, 2009; Hofmeister et al., 2013; Keller, 2000; Schütze, 1996; Sprouse, 2007; Sprouse et al., 2012; see Sprouse, 2023 for a review). But these sources are not visible in the acceptability judgment itself, as the process that generates the acceptability percept is assumed to combine the multidimensional error signal information into a single dimension. Linguistic theories are, of course, primarily interested in the source of the acceptability judgment, so that they can determine if effects are driven by the grammar (versus sentence processing), and if so, postulate constraints in the appropriate component of the grammar. Linguists deploy a number of different approaches to identifying the sources of acceptability effects, from manipulating known syntactic/semantic/sentence-processing effects within an acceptability judgment experiment, to attempting to correlate distinct data types with judgments. In this study, we would like to explore to what extent localization information (hemodynamic responses) gathered through fMRI can provide additional information for understanding the grammar and processing of constructions that are of central interest to (generative) syntacticians.

The idea to use fMRI to localize activation during acceptability judgments of violations is obviously not new. “Violation paradigms” have been in use in the fMRI literature since the early work in the 1990s (reviewed in more detail in section 2). The logic of violation paradigms is that there are (at least) two types of processes that fMRI might detect: the processes underlying the identification of the violation (i.e., a mismatch in grammatical requirements) and the processes deployed to repair the violation (i.e., to create an interpretable meaning from the utterance). The theoretical challenge, then, is to link these processes back to linguistic theories of the violations. But, in practice, this theoretical challenge has gone underexplored, because, in recent years, the results from violation paradigms have been attributed to general (or, at least, not linguistically- specific) processes like executive function and/or working memory (e.g., Kaan & Swaab, 2002; Novick et al., 2005, 2014; Rogalsky & Hickok, 2011). If true, this would suggest that the results of violation paradigms are less relevant to syntactic theory, because they do not reveal much about the grammatical source or sentence-processing dynamics of the violation. In light of this possibility, in recent years, fMRI researchers have moved away from violation paradigms, focusing instead on experimental designs that focus on grammatical sentences, often using “naturalistic” materials such as audiobooks. The results of these studies have been used to develop models of grammatical sentence processing, but with relatively loose connections to syntactic theories beyond the general type of grammar assumed by the models (see Brennan, 2016; Kandylaki & Bornkessel-Schlesewsky, 2019, for reviews). We’d like to reconsider this trend. We would argue that the fMRI results to violation paradigms could become more interpretable in two ways: (i) by testing multiple distinct violation types within the same experiment, such that overlapping activation could indicate shared processes, and non-overlapping activation could indicate distinct processes; and (ii) by comparing the fMRI results with other data types, like ERPs, to help circumscribe hypotheses about the potential processes that could be triggered by the violation, specifically identification and repair processes.

To that end, we investigate the fMRI responses to two violations thought to have distinct underlying grammatical sources as a first step in a larger research program: phrase-structure violations of the type in (1, with the illicit sequence underlined), and subject island violations of the type in (2, with the island structure indicated by brackets):

1. *Which candidate does the moderator of the panel think avoided the debate’s <u>about</u> <u>questions</u> healthcare?
2. **\***Which candidate does the moderator think [the speech by] ruined the debate’s questions about healthcare?

We chose these constructions because they likely have distinct underlying grammatical sources: subject islands involve long-distance dependencies, with several potential constraints that could give rise to the unacceptability (reviewed in section 2), while phrase-structure violations involve local-dependencies, and are generally thought to only be syntactic in nature. These two constructions likely lead to distinct sentence processing dynamics, particularly at the level of identification and repair of the violation. These are also two of the conditions investigated in Neville et al., (1991), which recorded EEG during an acceptability judgment task (reviewed in section 2), so we can potentially compare our localization results with the ERP results to further narrow the set of possible theories at both the level of grammar and sentence processing. (That said, the goal of the current study is not to directly localize the source of the ERP results, but rather to logically compare the distinct information that the three methods yield. fMRI is probably not the correct tool for investigating the cortical source of ERPs due to its limited temporal resolution; MEG may be a better tool, particularly if combined with concurrent EEG.)

Anticipating our results, we find that these two violation types activate (almost entirely) distinct cortical networks, with each network mapping to a previously identified processing network - semantic processing (Binder et al., 2009) for subject island violations and working memory (and reordering; L. Meyer & Friederici, 2016) for phrase-structure violations. We combine these results with the previous ERP results from Neville et al. 1991 to narrow the possible theories of the grammatical source and sentence processing dynamics of these violations, and to suggest future experiments that could help to probe these violations further. We also discuss how these results can be used as a starting point for a larger research program combining fMRI with acceptability judgments. Taking into account critical methodological considerations, and interpreting the results carefully, this approach might help to strengthen the link between syntactic theory and neurobiology (see also Grodzinsky & Friederici, 2006; Hagoort, 2016; Krauska & Lau, 2023; Matchin, 2023; Matchin & Hickok, 2020; Sprouse & Hornstein, 2016).

## 2. Background

In this section, we briefly we review some of the literature on violation studies in fMRI, theories of the source of phrase structure violations and subject island violations, and the ERP evidence from Neville et al. 1991 about the processes that might be triggered by these violations. The goal is not to exhaustively review these literatures (they are three distinct fields unto themselves), but rather highlight the information that we will use to potentially interpret the results of our fMRI study.

### 2.1 fMRI and syntactic violations

There is a large literature using neuroimaging methods to study syntactic processing. Though our study is specifically focused on violation paradigms, it is worth noting the broad array of studies on syntactic processing, as this provides part of the context for our study. Here we provide a sampling of citations for readers interested in exploring that context more deeply. For example, a number of studies have explored the central role of hierarchical structure (as opposed to linear sequential processing) in syntactic processing (e.g., Brennan et al., 2016; Ding et al., 2016; Matchin, İlkbaşaran, et al., 2022; Musso et al., 2003; Pallier et al., 2011), several have studied the processing of long-distance dependencies (e.g., Just et al., 1996; Stromswold et al., 1996, Makuuchi et al., 2013; Santi et al., 2015), and many have compared syntactically licit sentences or phrases with unstructured word lists (e.g., Bemis & Pylkkanen, 2011; Matchin et al., 2017; Mazoyer et al., 1993; Pallier et al., 2011; Rogalsky & Hickok, 2009; Zaccarella & Friederici, 2015). Though these studies have been critical in developing theories of the neurobiology of grammatical syntactic processing, the connection with syntactic theory has often been relatively tenuous, perhaps because syntactic theory is typically concerned with ungrammatical sentences that can be used to identify constraints in the grammar.

A more direct link between the two literatures could be facilitated by fMRI studies of violations. And to date, there have been a number of such studies, including studies of phrase structure violations of different types (Embick et al., 2000; Friederici et al., 2003; Kuperberg et al., 2000, Meyer et al., 2000; Moro et al., 2001; Newman et al., 2001; Rüschemeyer et al., 2005), tense violations (Kuperberg, Holcomb, et al., 2003; Ni et al., 2000), and agreement violations (Folia et al., 2009; Husband et al., 2011; M. Meyer et al., 2000). These violation paradigm studies have generally elicited activation in language-related regions such as left IFG (Broca’s area) and various portions of the left temporal lobe (see Hagoort & Indefrey, 2014 for a meta-analysis)). Because these regions have also been shown to support executive function resources such as cognitive control and working memory, some researchers have argued that the responses obtained in the violation paradigm studies do not reflect specific information about the grammatical source of the violations or about the sentence processing dynamics of the violation (Kaan & Swaab, 2002; Novick et al., 2005, 2014; Rogalsky & Hickok, 2011).

In order to address the concerns that violations could induce increased activation in executive function systems, we decided to contrast two distinct violation types, involving different potential syntactic constraints. Our logic is to triangulate responses specific to the particular syntactic mechanisms themselves by ascertaining the extent to which the activation maps for phrase structure violations and subject islands overlap. Regions activated selectively by one violation type and not the other would suggest specificity to the particular identification or repair processes involved in that type. Regions activated by both violation types would suggest a generic violation response, which we might expect in executive function regions. Phrase-structure violations have been relatively well studied within fMRI, so our goal was to replicate the previously observed activation pattern: activation in LIFG and left STG/MTG. To our knowledge, subject islands have not been studied using fMRI (Christensen et al., 2013 did study wh-islands in Danish, but crucially, it has been claimed that Danish does not have a wh-island constraint in the grammar starting from at least Erteschik-Shir, 1973, which was corroborated by the results in their study). Therefore, the pattern that we observe will be a completely novel contribution to the localization of violations, and potentially reveal new pathways for interpreting violation results.

### 2.2 Theories of the sources of the violations, and predictions for identification and repair processes

For this study, we selected two syntactic violation types that are relatively distinct: phrase structure violations and subject island violations. The specific phrase structure violation that we chose (illustrated in 1 above) is created by taking the sequence possessive noun + noun + preposition, and transposing the noun and preposition. Prepositions cannot follow possessive nouns in English, so this is a violation of the syntactic selectional requirements of possessive nouns. To our knowledge, this kind of violation is treated relatively uniformly across different theories of syntax – it is a violation of the local dependency between the possessive noun and the noun that follows it. We selected it because there is no uncertainty around the syntactic analysis of this violation and because it has been studied extensively in the fMRI and EEG literatures.

The assumption in those literatures is that the identification processes are likely related to local (syntactic) predictive mechanisms, and that the repair process is likely working memory, as the parser can search through potential permutations of the immediately adjacent items for a grammatically licit sequence. In this way, phrase structure violations can serve as a relatively stable comparison point for exploring the effects of different syntactic violation types.

We selected subject islands (illustrated in 2 above) as a comparison because (i) they instantiate a long-distance dependency violation (rather than a local dependency violation), (ii) they have been studied together with phrase structure violations in an influential EEG experiment (Neville et al., 1991), and, most importantly from a grammatical point of view, (iii) there are at least three distinct proposals in the literature for the source of subject islands – syntax-based theories, information-structure-based theories, and processing-based theories.

Syntax-based theories define special syntactic domains that are opaque to the syntactic operation (e.g., movement) that creates the long-distance dependency. There are different approaches to this in the syntax literature that could potentially explain subject island violations, such as Subjacency (e.g., Chomsky, 1986), the Condition on Extraction Domains (e.g., Huang, 1982), Phase Impenetrability (see Boeckx, 2013 and Citko, 2014 for reviews), and Multiple Spell-Out (Uriagereka et al., 1999). Information-structure-based approaches analyze island effects as a clash between the focus created by wh-movement and the backgroundedness of the island constituents that the wh-word is extracted from (Abeillé et al., 2020; Chaves & Dery, 2019; Erteschik-Shir, 1973; Goldberg, 2006). Subject islands would then arise if the information contained in the subject were considered backgrounded in English. Finally processing-based theories explain island effects through a limitation in the resources available to successfully process both the long-distance dependency and the constituent that we call an island. For example, Kluender & Kutas (1993b) propose a limitation in working memory capacity (see Kluender, 2004 specifically for subject islands), while Deane (1991) proposes a limitation in attentional resources. Because fMRI results tend to be classified in relatively broad terms, for this first project, we will abstract away from the detailed mechanisms of individual theories, and instead consider them as classes of theories: syntactic, information-structure, and processing.

The three classes of island theories might be plausibly interpreted to make distinct predictions about the identification and repair processes triggered by island violations. For syntactic theories, the identification processes would likely be related to long-distance dependency processing and gap identification, perhaps implicating working memory retrieval and predictive processing mechanisms. The repair processes would likely be related to the thematic structure of the sentence, as the displaced filler cannot be integrated into the thematic structure of the sentence and the prepositional phrase within the subject is missing a thematically-required argument. For information-structure theories, both the identification and repair strategies would be related to the focus structure of the sentence, as the focused-nature of the wh-dependency clashes with the backgrounded nature of the prepositional phrase in the subject. Finally, for processing theories, the identification processes would be related to recognizing an overload of working memory capacity, and the repair processes would be the strategic reduction or reallocation of working memory resources (e.g., the reduction process suggested by Kluender & Kutas, 1993a). These qualitative distinctions among the processes predicted by the three theories might plausibly arise as distinct patterns of activation in fMRI.

### 2.3 Previous ERP results

Though our goal is not to investigate (or localize) specific ERPs, we do wish to take advantage of the fact that islands and syntactic violations more broadly have been studied more extensively in the ERP literature. Because we are using a different method, fMRI, to examine the spatial dimensions of these violation effects, it is important to first consider the relationship between the neurophysiological signature of ERP effects using EEG and the hemodynamic response of fMRI. fMRI BOLD signal is correlated at the neurophysiological level by local field potentials, as captured by EEG, which reflect high-frequency synchronous dendritic activity across large populations of neurons (Logothetis, 2008; Logothetis et al., 2001). However, studies of ERPs (using intracranial EEG) and fMRI in separate, parallel experiments with different participants, have shown substantial divergences in the functional response across brain regions (Engell et al., 2012; Huettel, 2004). Thus, while in many cases there appear to be converging effects across ERPs and fMRI (e.g. Lau et al., 2008; Linden, 1999; Matchin et al., 2019; Stevens et al., 2000), caution is warranted in making straightforward predictions for one modality based on the other.

A particularly important previous ERP study is Neville et al., 1991, which studied both phrase structure and subject island violations together in the same experiment. For phrase structure violations, Neville et al. (1991) found an ELAN, LAN, and a P600; and for subject island violations, they found an increase in P2 amplitude and a P600. We will set aside the ELAN because of recent work suggesting it may be an artifact that arises when comparing two conditions that do not match in the word preceding the critical word (Steinhauer & Drury, 2012). But the other ERPs provide potential evidence for the kinds of processes that are triggered by these violations. Here we provide a brief review of possible functional interpretations for the effects that Neville et al. observed.

The LAN is elicited by two types of constructions that appear, at least superficially, to be distinct: morphosyntactic violations (e.g., Coulson et al., 1998; Friederici et al., 1996; Münte & Heinze, 1994) and grammatical sentences that likely require higher working memory resources such as long-distance dependencies (e.g., Kluender & Kutas, 1993a) and garden-path sentences (Kaan & Swaab, 2003). The primary challenge in the LAN literature is explaining this apparent dichotomy. One approach is to simply conclude that there are two distinct sources for the LAN, one for morphosyntactic violation detection and one for working memory processes (e.g., Martín-Loeches et al., 2005; Molinaro et al., 2011). The second approach is to postulate that the only source is working-memory processes (as the LAN arises independently in working-memory tasks). One way to explain morphosyntactic LANs under a working memory theory is to identify working-memory processes that could be triggered by the violation detection process or by attempts to repair the violation. For phrase structure violations, a repair process that reorders items from memory is a likely candidate. A second way to explain morphosyntactic LANs under a working memory theory is to postulate that they are not really LANs, but rather an artifact of averaging across two groups of participants – one group that shows an N400 to the violation, and one group that shows a P600 to the violation. Tanner & Van Hell (2014) argue extensively for this latter approach, noting that it also explains why morphosyntactic LANs typically only appear together with P600s. An N400 seems unlikely for phrase structure violations, but we note it as a logical possibility.

To our knowledge, the P2 has not been extensively studied within the sentence processing or syntactic violation literature. Instead, the P2 has primarily been investigated within the lexical processing literature, where the P2 has been linked to prediction/expectation mismatches (see Almeida & Poeppel, 2013 for some discussion). Given that, and given that Neville et al. (1991) do not provide a functional interpretation of the P2 in their discussion, for this study we will tentatively interpret the P2 as indexing prediction/expectation mismatches, and note that there is need for future work investigating the functional interpretation of the P2 in sentence processing.

The P600, like the LAN, is elicited by a wide range of conditions, such as morphosyntactic violations (e.g., Friederici et al., 1993; Hagoort et al., 1993; Osterhout & Holcomb, 1992), garden-paths (e.g., Friederici et al., 1996; Kaan & Swaab, 2003; Osterhout et al., 1994), long-distance dependency resolution (e.g., Fiebach et al., 2002; Kaan et al., 2000), and unexpected thematic role assignments (e.g., Kim & Osterhout, 2005; Kuperberg, Sitnikova, et al., 2003). Perhaps unsurprisingly, the range of conditions that elicit P600s has led to a number of proposals for its functional interpretation. Here we will mention three. One approach is to analyze the P600 as indexing a very general structure-building process, such as *unification* in Hagoort’s (2005) memory-unification-control model, which is used to build phonological, syntactic, and semantic representations, or *integration* in Brouwer et al.’s (2012) retrieval-integration model, which is used to integrate material into the “mental representation of what is being communicated.” A second possibility is that the P600 reflects reanalysis or repair processes (as argued by Neville et al. 1991 for subject islands), which would explain P600s to both morphosyntactic violations and garden-path sentences, but perhaps not long-distance dependency resolution (which would still require a memory-retrieval or integration analysis). A final possibility is that the P600 is a temporally-delayed version of the P3b, which is known to track mismatches in expectations (e.g., Coulson et al., 1998; Osterhout, 1999). Again, this would potentially explain morphosyntactic violations and garden-path sentences, but perhaps not long-distance dependency resolution.

### 2.4 The logic of combining this information

With this review in hand, we can better articulate the logical inference process we will follow here. First, we will look at the activation patterns that we observe and compare them to previous results in the fMRI literature to identify potential functional interpretations. Next, we will look for overlap in the activation patterns of the two violations, which we will take to be indicative of shared processes (perhaps general processes like executive control, general error detection, or general repair), and for non-overlap, which we will take to be indicative of processes that are unique to each violation. Finally, we will use the information reviewed above about the potential identification and repair processes triggered by encountering the violations to draw potential inferences about the grammatical source of the violations, although our conclusions are limited.

## 3. Materials & Methods

### 3.1 Participants

We recruited 17 right-handed, adult native speakers of English for this study. Sample size was determined based on previous studies using similar sample sizes. Participants had normal or corrected-to-normal vision, no hearing impairment, and reported no history of neurological disorder. Participants were paid $30 an hour for their participation. Consent was acquired from each participant before the experiment and all procedures were approved by the Institutional Review Board of UC Irvine.

### 3.2 Stimuli

120 grammatical sentences (Grammatical), e.g. sentence (3), were generated with the following identical structure: a WH-question, with a matrix clause subject with a prepositional phrase modifier, the matrix verb *think*, an embedded clause as the complement of *think*, with the embedded subject extracted to the initial position of the entire sentence to form the WH-question. The embedded clause contained a transitive verb with a noun phrase object consisting of a possessive noun, signaled by*‘s*, and a noun phrase consisting of a noun with a prepositional phrase modifier. A subject island version of each of these grammatical sentences (Subject Island), e.g. sentence (4), was generated by removing the prepositional phrase modifier from the matrix subject, adding it to the embedded subject, and extracting the object of the prepositional phrase in the embedded subject, which is recognizable as a subject island structure. A phrase structure violation version of each of the grammatical sentences (PSV), e.g. sentence (5), was generated by inverting the order of the preposition and noun in the embedded object. Finally, a combined violation version of each of the grammatical sentences (Both), e.g. sentence (6), was created by combining the modifications for subject islands and PSV. All the materials created for this experiment are available in the Appendix.

1. **Grammatical:** *Which candidate* does the moderator of the panel think avoided the debate’s questions about healthcare?
2. **Island:** \**Which candidate* does the moderator think [the speech by] ruined the debate’s questions about healthcare?
3. **PSV:** \**Which candidate* does the moderator of the panel think avoided the debate’s <u>about questions</u> healthcare?
4. **Both:** \**Which candidate* does the moderator think [the speech by] ruined the debate’s <u>about questions</u> healthcare?

The “Both” condition is primarily exploratory (completing a 2x2 design). However, we note that it also allows us to maximize power, as we can measure the main effects of subject island and PSV by combining data across all conditions.

We included one additional condition: subvocal articulatory rehearsal. We included this condition in order to isolate brain networks associated with verbal working memory, as articulatory rehearsal substantially overlaps networks associated with verbal working memory tasks (Hickok et al., 2003) and this task has been used previously in fMRI studies of sentence processing (Matchin et al., 2014; Rogalsky et al., 2008, 2015). However, we only report limited results of the analysis of the articulatory rehearsal condition because the extent and magnitude of brain activations associated with this condition were very high relative to our sentence violation contrasts of main interest. Activations for the rehearsal condition completely enveloped both of the violation effects and thus have extremely limited inferential value in this context.

We created 120 lexically matched quadruplets instantiating the four conditions, which were matched for both lexical content and number of syllables across conditions. We distributed the items into 4 lists, each containing 30 tokens per condition, using a Latin Square procedure. The stimuli were recorded by a phonetically trained native speaker of American English in a sound-attenuated booth. The materials were down-sampled from 44,100 to 22,050 Hz. All stimuli were equated for mean root-mean square power, intensity, and controlled for duration across conditions and across lists: Grammatical average duration 4.911 (SD 0.360), PSV average duration 4.923 (SD 0.390), Island average duration 4.910 (SD 0.571), Both average duration 9.24 (SD 0.363); List 1 average duration 4.933 (SD 0.367), List 2 average duration 4.873 (SD 0.564), List 3 average duration 4.932 (SD 0.377), List 4 average duration 4.932 (SD 0.377). All editing was done on Praat software (Boersma & Weenink, 2011).

### 3.3 Experimental Procedure

The speech stimuli were presented over eight scanning runs with an average length of about five minutes per run, plus a high-resolution anatomical MRI collected at the end of the experiment, thus each participant was in the scanner for almost an hour. Each run contained 30 sentences, which were on average 4.92 seconds. The order of sentences for each list was randomized across runs, such that there were variable numbers of trials in each condition presented per run, but equated across runs. Each subject was presented with two different stimulus lists in order to yield 60 trials per condition per participant. The interstimulus interval between sentences was fixed at 2500 ms. The order of presentation of runs was randomized for each participant. Stimulus delivery was performed using Cogent 2000 software for MATLAB (MathWorks, Inc.) with MRI-compatible electrostatic headphones. Participants were told that they would hear utterances in English. Participants were asked to provide a binary forced-choice acceptability judgment for each sentence (with the options of acceptable or unacceptable). Participants were also instructed to listen carefully for meaning, as they would have comprehension questions to answer about some of the sentences at the end of each run. At the end of each run, four true/false comprehension questions about four of the sentences were presented. Half of the questions were true; half were false. Throughout stimulus presentation, a black fixation cross appeared in the center of the screen. Each of the four functional scanning runs included one 30-second period of subvocal articulatory rehearsal, randomly cued at some point during the run. Participants were instructed to repeat the sequence /ba/-/da/-/ga/ as rapidly as possible when the fixation cross flashed between red and blue. In addition, there was a random 30s period of rest without stimulation once per run, in order to facilitate with baseline estimation.

### 3.4 fMRI data collection, processing, and analysis

MR images were obtained in a Philips Achieva 3T (Philips Medical Systems, Andover, MA, USA) fitted with an eight channel RF receiver head coil at the high field scanning facility at UC Irvine. For 15/17 participants, we first collected a total of 1200 T2*-weighted EPI volumes over 8 runs using Fast Echo EPI in ascending order (TR = 2 s, TE = 30 ms, flip angle = 90**°**, in-plane resolution = 1.85 mm × 1.85 mm, slice thickness = 5 mm with 0.5 mm gap). Two participants were unable to complete one run of the experiment, so 1050 volumes over 7 runs were collected. Following the experiment, a high-resolution T1-weighted anatomical image was acquired in the axial plane (TR = 11 ms, voxel size = 1 mm isotropic).

Data preprocessing was performed using AFNI software (Cox, 1996; http://afni.nimh.nih.gov/afni). The first five volumes of each run were discarded to control for T1 saturation effects, followed by slice-timing correction. Motion correction was achieved by using a 6-parameter rigid-body transformation, with each functional volume in each run first aligned to a single volume in that run. Functional volumes were aligned to the anatomical image, and subsequently aligned to the Talairach template brain (Talairach and Tournoux, 1988). Functional images were resampled to 3 mm isotropic voxels, spatially smoothed using a Gaussian kernel of 6 mm FWHM, and converted to percent signal change values within each run.

First-level (single subject) analyses were performed on each individual’s data using AFNI’s 3dDeconvolve function. The regression analysis was performed to find parameter estimates that best explained variability in the data. For all four sentence conditions, each predictor variable representing the timecourse of stimulus presentation was entered into a deconvolution analysis that estimated parameters best representing the timecourse of the hemodynamic response function. For the blocked articulatory rehearsal condition, the stimulus timing regressor was convolved with a canonical hemodynamic response function. The six motion parameters were included as regressors of no interest. The data were detrended for low frequency noise at the first-level analysis stage using the ‘polort’ parameter with a value of 2 (linear and quadratic trends).

Second-level (group) analyses were performed by entering the parameter estimates for percent signal change for each condition for each participant into AFNI’s 3dANOVA2 function. We analyzed the following effects: each individual sentence condition and articulatory rehearsal vs. rest, the main effect of SUBJECT ISLAND (subject island + both > PSV + good), the main effect of PSV (PSV + both > subject island + good), and the interaction. To correct for multiple comparisons across voxels, we used a family-wise error (FWE) cluster size correction using Monte Carlo simulations. First, we estimated the spatial autocorrelation function in each ’s data using AFNI’s 3dFWHMx function with the *acf* option (Cox et al., 2017). We then averaged the parameter estimates across s and ran cluster simulations to estimate the Type 1 error rate using AFNI’s 3dClustSim and the *acf* option. We adopted a voxel-wise statistical threshold of *p* < .005 (one-tailed) and a cluster extent threshold of 77 voxels (2079 mm^3^) to keep the FWE rate at *p* < .05. In order to identify the broader networks involved in each violation type, we reduced the voxel-wise threshold to p < 0.01 (one-tailed) and used a cluster extent threshold of 20 voxels. Using a looser threshold might help to identify a broader network, rather than an isolated region, strengthening our confidence in the previously established functionality of that network that would be difficult if we only identified a small number of individual regions. This would facilitate making reverse inferences based on the results, which depends critically on the confidence of the functionality of the brain networks identifed (Poldrack, 2006). We then created overlap maps among the main effects of SUBJECT ISLAND and PSV.

Although our primary analyses were performed at the group level, we note that group analyses may obscure the true nature of underlying effects that exist at the individual participant level (Fedorenko & Kanwisher, 2009). Therefore, in order to bolster the conclusions we draw from the group-level activation maps, we also report activation maps for each individual participant for the same main effects of interest described above: the main effects of SUBJECT ISLAND (subject island + both > PSV + good) and PSV (PSV + both > subject island + good), using an uncorrected one-tailed voxel-wise threshold of p < 0.001.

## 4. Results

### 4.1 Behavioral results

The top panel of figure 1 reports the results of the binary forced-choice acceptability judgment task collected during scanning. Participant responses were coded as 1 (acceptable) or 0 (unacceptable). The bar plot is a histogram of the by-participant means; the vertical blue line is the grand mean of all participants. The Grammatical, Phrase Structure Violation, and Both conditions behaved as expected: Grammatical has a relatively high rate of being categorized as acceptable, and PSV and Both have relatively low rates. But the rating for Subject Islands was surprising, as the grand mean is near the middle of the range (0.61), and about half of our participants rated subject islands acceptable more than 80% of the time. This is surprising because subject island violations using the same structure are reliably rated relatively low in acceptability tasks across a number of published studies (e.g., Sprouse et al., 2011, 2012, 2016), therefore we expected these to be characterized as unacceptable more frequently.

**Figure 1:**
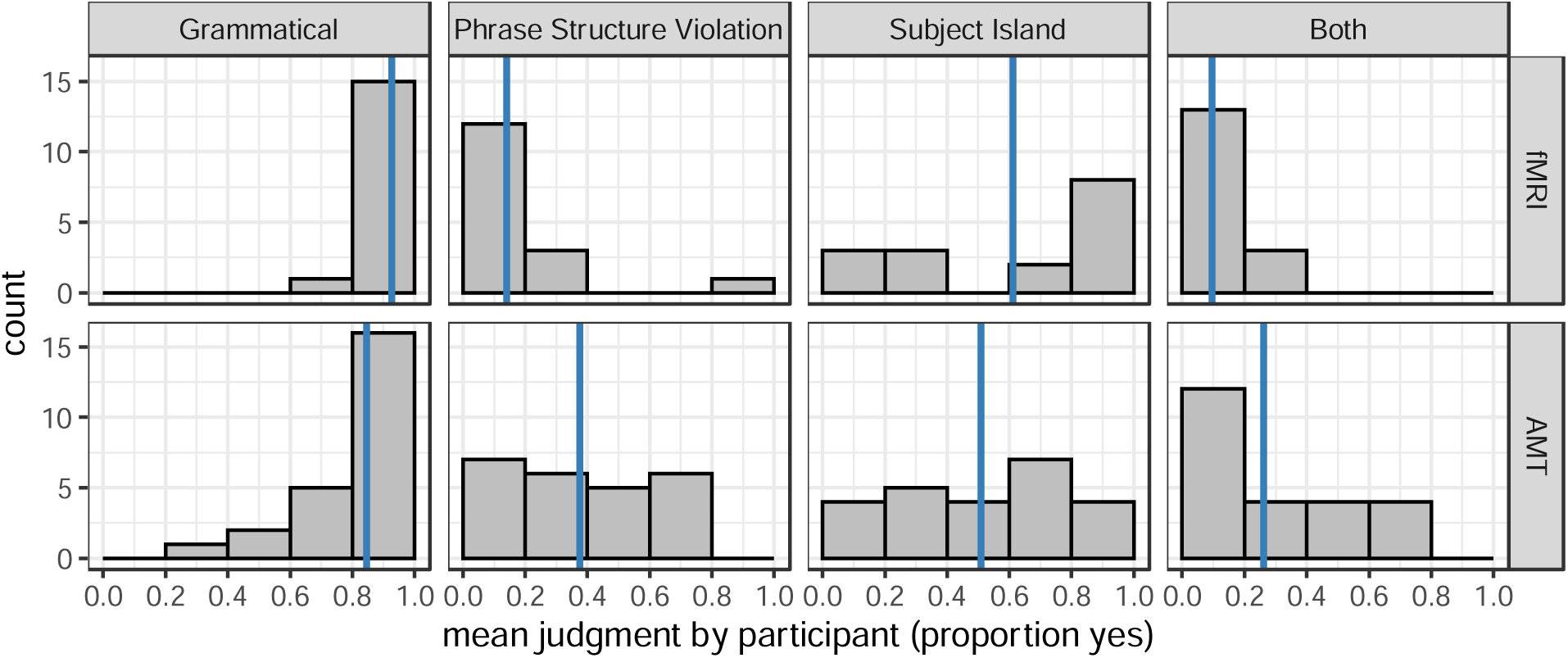
Histograms of the by-participant mean acceptability ratings for the judgments collected during fMRI scanning (top row) and a shortened replication using Amazon Mechanical Turk (bottom row; AMT). The blue line in each plot represents the mean rating for that condition.

Our first suspicion was that perhaps the increased noise during scanning made it difficult for participants to parse the subject island violations, and they defaulted to a higher rating, perhaps because the violation itself is not easy to characterize, and therefore not easy to correct with a simple change to the word order (see Crain & Fodor, 1987 for arguments that “correctible” violations and “uncorrectible” violations behave differently in psycholinguistic tasks). To test this hypothesis, we ran an acceptability judgment experiment on Amazon Mechanical Turk (AMT) using the same materials (the same auditory recordings), the same task (binary forced-choice), and a parallel-but-shortened survey design – 12 tokens per condition rather than 60. We recruited 24 self-reported native speakers of English (a similar sample size to the 17 in the fMRI study). The results of this experiment are in the bottom row of figure 1, and crucially, are very similar to the results collected during fMRI scanning: Grammatical is rated relatively high, PSV and Both are rated relatively low, and Subject Islands are rated near the middle of the scale. It does not seem to be the noise of the scanner that drove the results.

Our second suspicion is that this could be a satiation effect, that is, an increase in acceptability after repeated exposure to a given violation type. Though the results in the satiation literature are inconsistent (see Chaves & Putnam, 2020 for a review), we nonetheless looked for satiation in both the fMRI sample and the AMT sample. Figure 2 plots the mean rating of the Subject Island condition by exposure (1 through 60 for the fMRI sample; 1 through 12 for the AMT sample). As is clear in the figure, there is no obvious sign of satiation. To statistically corroborate this, we created linear mixed effects models with acceptability as the dependent variable, exposure order as a fixed effect, and participant as a random effect (intercept only). We calculated *p*-values for the fixed effect of exposure order using the lmerTest package (Kuznetsova et al., 2017), which uses the Satterthwaite approximation for degrees of freedom to derive an F test from the linear mixed effects model. For the fMRI sample, we created models that looked at every exposure individually, every two exposures, every 5 exposures, every 10 exposures, and splitting the experiment in half (30 exposures). None were significant: *p*=.80, *p*=.81, *p*=.79, *p*=.99, and *p*=.87, respectively. For the AMT sample, we created models for every exposure, every two exposures, and half the experiment (6 exposures). Again, none were significant: *p*=.81, *p*=.96, and *p*=.69, respectively. It does not seem that the surprisingly high ratings for the subject island violations were due to radiation.

**Figure 2:**
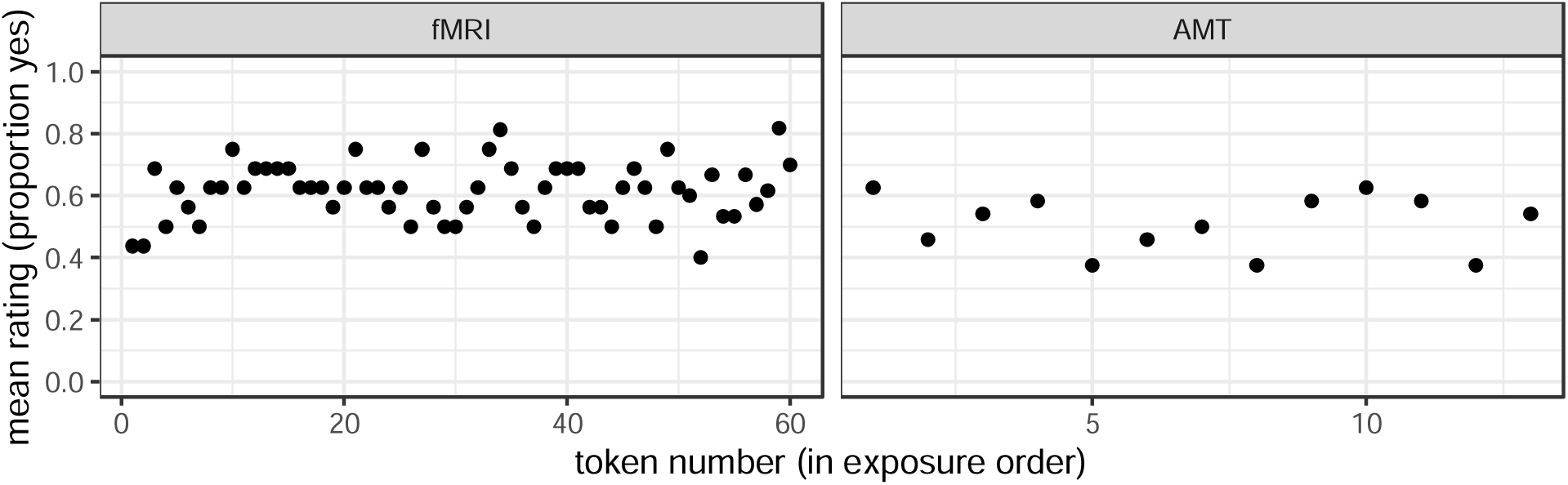
The mean acceptability rating of Subject Island Violations by exposure order in the two acceptability judgment experiments experiments.

Our post-hoc hypothesis is that the surprisingly high ratings for the subject island violations are due to an interaction between the binary nature of the task and the fact that 75% of the items in our experiment (both the fMRI sample and the AMT sample) are ungrammatical. If participants expect a relatively even distribution of the two response categories in the task, then participants could decide to shift one of the conditions into the “acceptable” category. Of the three unacceptable conditions, the subject island violation may be the most likely to be recategorized as acceptable given that the violation is not easy to characterize compared to the phrase structure violation in the other two conditions (again, see Crain & Fodor, 1987). Testing this hypothesis is far beyond the scope of this study, as it would require systematically testing different acceptability tasks, different ratios of unacceptable to acceptable items in an experiment, and different types of violations. Fortunately, we can use the fMRI data to determine if the Subject Islands were truly processed as if they were acceptable sentences: if the fMRI results for Subject Islands shows no difference compared to the Grammatical condition, then we can conclude that they were truly processed as acceptable; if there is a difference, we can conclude that the more frequent than expected categorization as “acceptable” was a superficial task effect. That said, we will consider this issue in the interpretation of our fMRI relative to the Neville et al. ERP results in Section 5.4, as the ratio of acceptable to unacceptable items has been shown to affect P600s.

4.2 fMRI results

Figure 3 displays the results of the main effects of each violation type for our group anlayses. The main effect of PSV identified one significant cluster: the left posterior inferior temporal gyrus (center of mass: -49, -47, -12, 106 voxels). The main effect of SUBJECT ISLAND identified two significant clusters: one in the left ventrolateral prefrontal cortex (VLPFC) (center of mass: -35, 45, 5, 109 voxels), and one in the left dorsomedial prefrontal cortex (DMPFC) (center of mass: - 32, 17, 46, 80 voxels). The interaction analysis did not yield any significant clusters; the only cluster that approached significance was found in the right inferior precentral and postcentral gyrus, indicating that none of our main effects were being driven predominantly by one condition.

**Figure 3.**
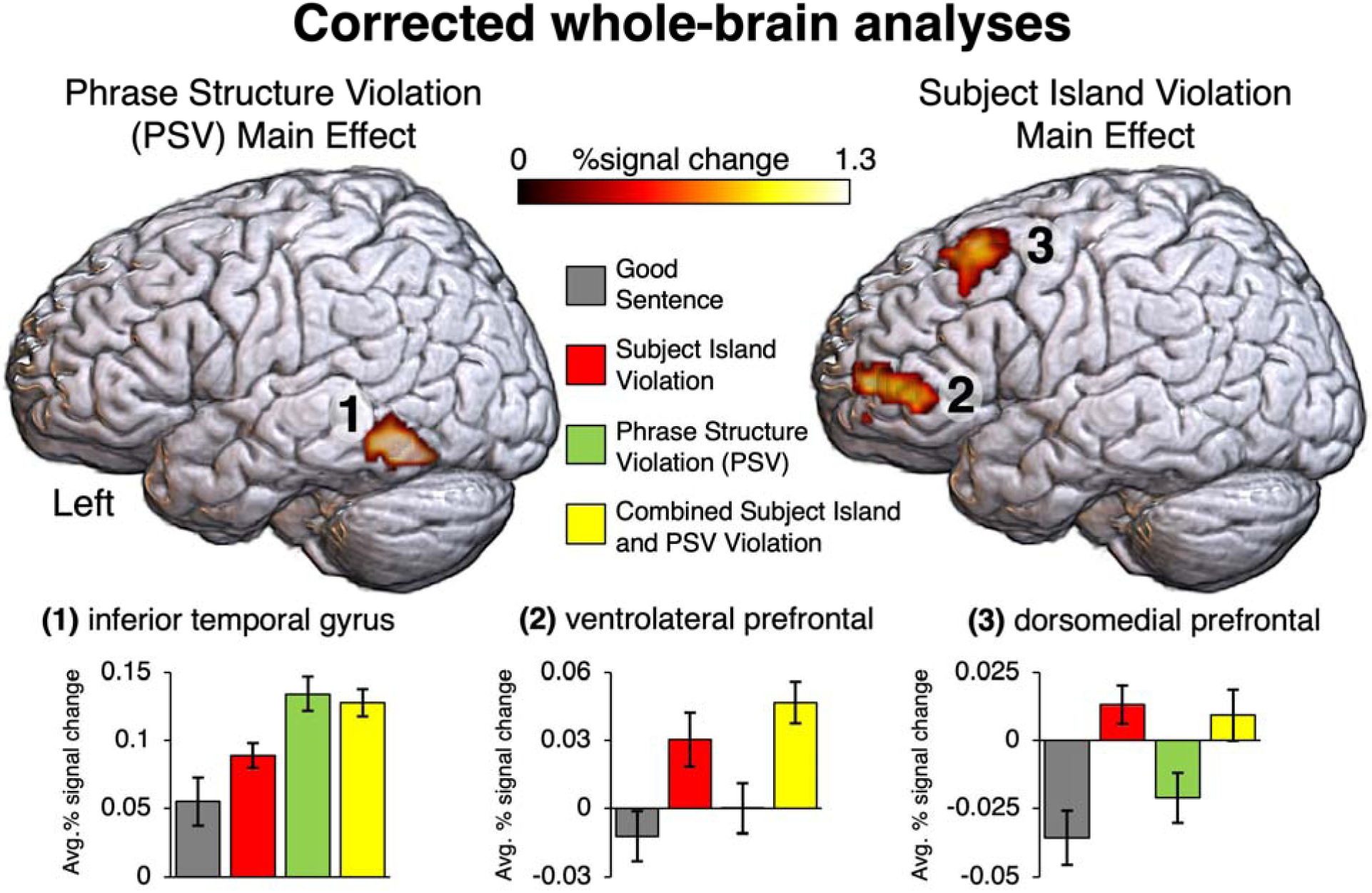
TOP: Significant clusters for the main effect of phrase structure violation (PSV) (left) and Subject Islands (right). BOTTOM: extracted percent signal change averaged within each significant cluster for each individual sentence condition. Error bars depict standard error with subject effects removed (Cousineau, 2005). NOTE: statistics are calculated at the voxel-level, corrected for multiple comparisons using cluster size simulations. Bar plots illustrating effect sizes within each significant cluster are for visualization of results, not for statistical inference.

The uncorrected, reduced threshold group analyses (voxel-wise p < 0.01) identified two largely distinct, non-overlapping networks for the main effects of SUBJECT ISLAND and PSV (Figure 4). In addition to the significant effects in VLPFC and DLPFC, the main effect of SUBJECT ISLAND activated the left and right anterior temporal lobe, posterior middle temporal gyrus, and angular gyrus. By contrast, in addition to the significant main effect of PSV in the left inferior temporal gyrus, this contrast activated left posterior superior temporal gyrus (pSTG), left posterior superior temporal sulcus (pSTS), left posterior middle temporal gyrus (pMTG), left posterior IFG (pars opercularis), and bilateral anterior inferior parietal lobe. There was very little overlap between the increased activation for SUBJECT ISLAND and PSV at this reduced threshold (5 voxels total), centered on the left pSTS/MTG.

**Figure 4.**
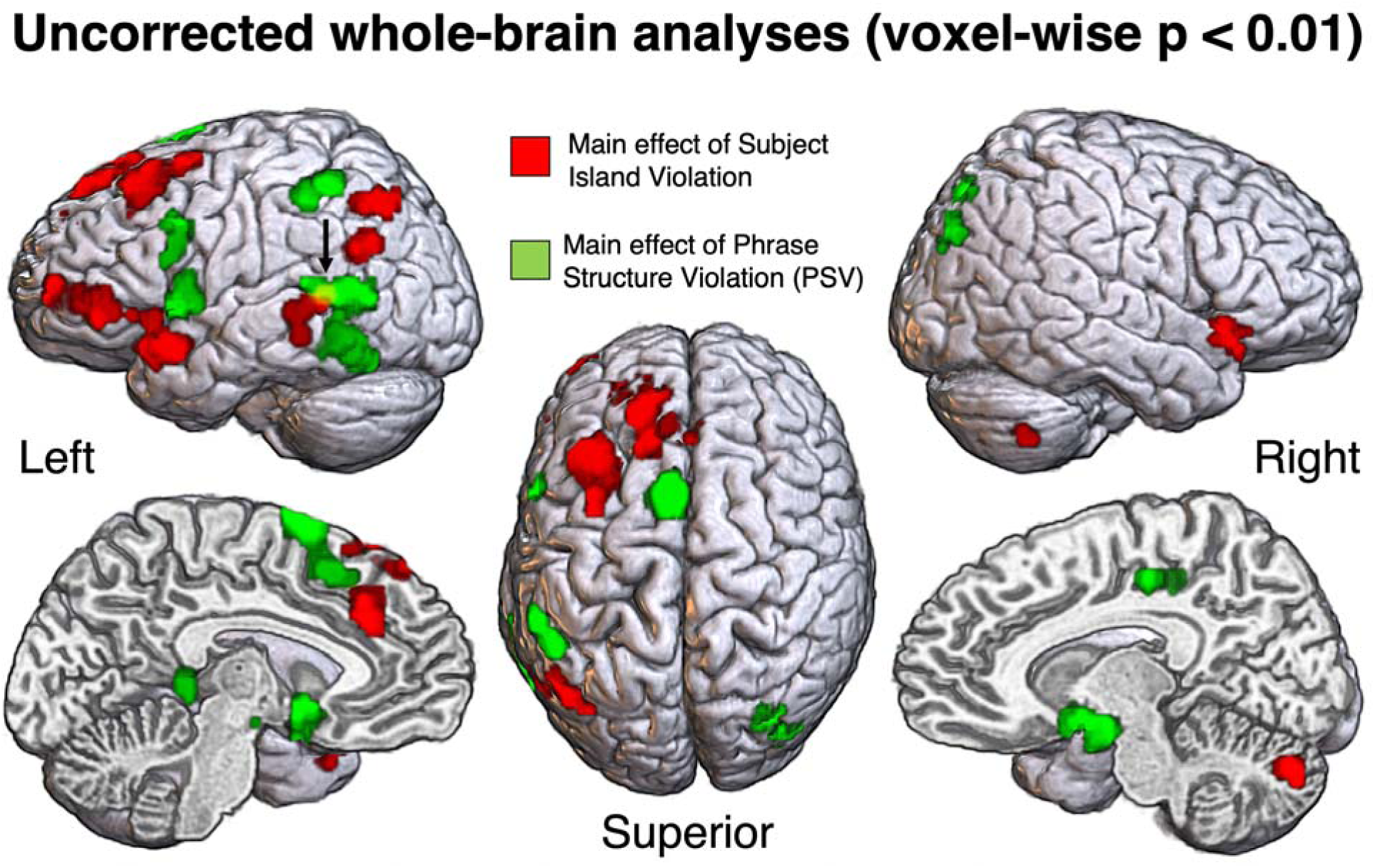
Overlap maps for the reduced threshold analyses (voxel-wise p < 0.01) for Subject Island and Phrase Structure Violations (PSV). RED: main effect of Subject Island Violation, GREEN: main effect of PSV, YELLOW: overlap. The arrow points towards the small overlap between these two main effects (5 voxels).

The rehearsal condition activated a robust, bilateral network involving posterior temporal lobe (involving superior, middle, and inferior temporal gyri), anterior inferior and superior parietal lobe, anterior insula, and frontal lobe spanning the precentral, inferior, and middle frontal gyri. The only areas with very minimal or no activation in the rehearsal condition included the bilateral anterior temporal lobes and posterior, inferior parietal lobe as well as middle-anterior superior frontal gyrus. As discussed above, we limited our discussion of these effects given the limited inferential value such widespread and robust activations for this condition allow.

### 4.3 Individual participant fMRI results

The uncorrected individual participant activation maps (voxel-wise p < 0.001) (Figure 5) showed that the main effects of SUBJECT ISLAND and PSV were substantially variable across participants, both in terms of the spatial distribution of these effects as well as overall statistical robustness.

**Figure 5.**
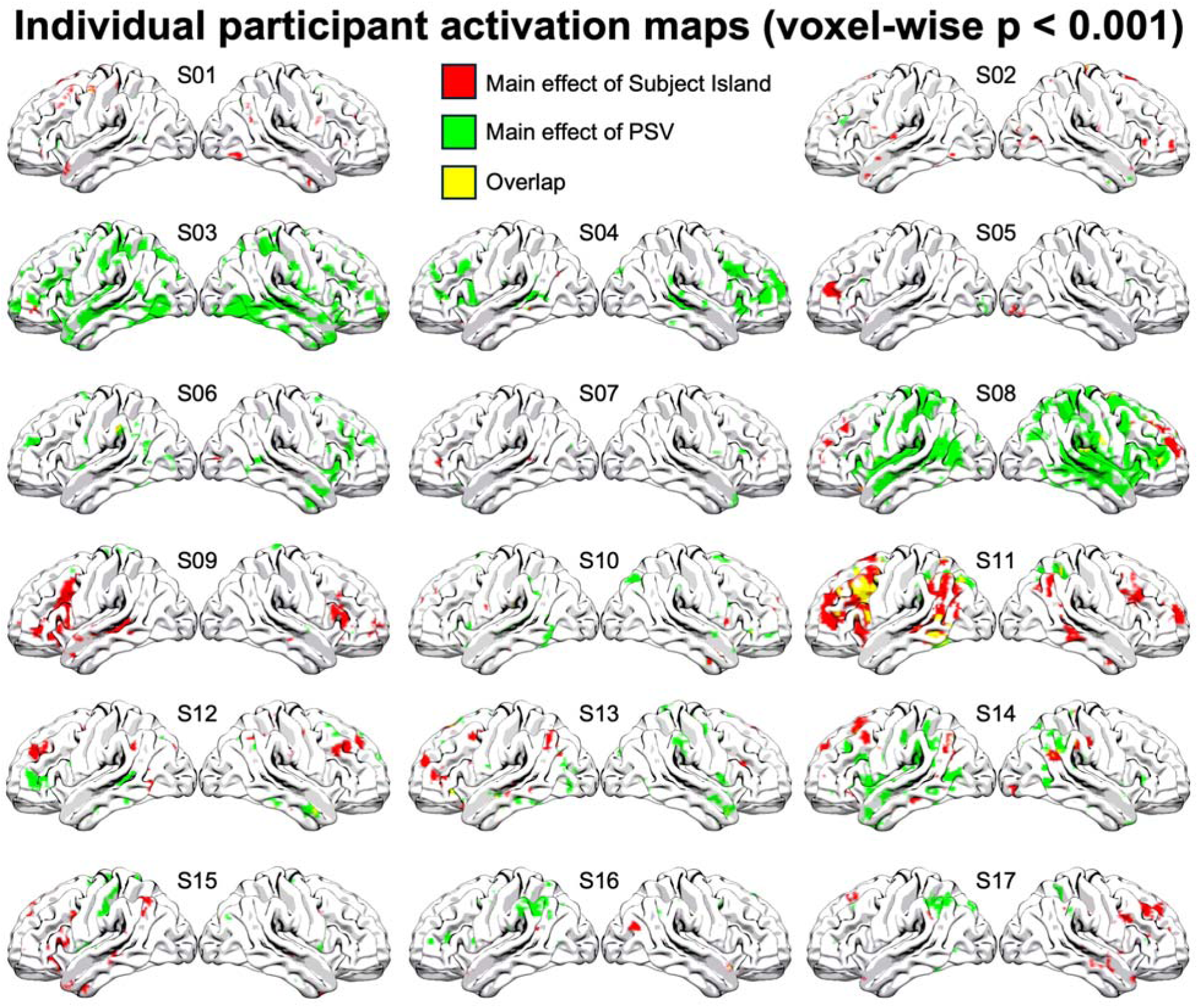
Individual participant activation maps (voxel-wise p < 0.001) for the main effects of Phrase Structure Violations (PSV) and Subject Island violations, displayed on an inflated cortical surface map for maximum visbility. RED: main effect of Subject Island Violation, GREEN: main effect of PSV, YELLOW: overlap.

Three participants (S03, S06, S10) showed notable activation for PSV and not for Subject Islands, two participants (S05, S09) showed notable activation for Subject Islands and not for PSV, and three participants (S01, S02, S07) showed minimal effects for either violation type. The remaining nine participants (S04, S08, S11, S12, S13, S14, S15, S16, S17) did show substantial activation for both of the violation types. Of these participants, there was very limited overlap for these two main effects, with the exception of participant S11, for whom there was significant overlap of these effects in posterior middle-inferior temporal lobe, inferior parietal lobe, and posterior inferior-middle frontal cortex. Certain tentative trends in the individual maps that contributed to the group-level activation maps were observable, including activation for PSV in posterior inferior temporal lobe, and activation in dorsomedial and ventrolateral prefrontal cortex, inferior angular gyrus for Subject Islands.

## 5. Discussion

### 5.1 The activation of distinct cortical networks for each violation type

Given that both subject islands and phrase structure violations (PSV) deviate from grammatical sentence structures and involve degraded acceptability, one might have expected substantial overlap between these conditions in brain networks involved in executive function (to detect the violations, and perhaps respond to the processing demands of any repairs). However, PSV and subject islands produced almost entirely non-overlapping activation maps. Both violation types elicited activation in prefrontal cortex, but in distinct regions. The responses in prefrontal cortex might reflect executive functions such as cognitive control or working memory needed to detect the violation and repair the structure, but the prediction that violations of different types would produce a common generic violation response was not upheld. The only region that showed any overlap was in the left posterior superior temporal sulcus/middle temporal gyrus, a brain region long thought to underlie lexical processing, (Friederici, 2017; Hickok & Poeppel, 2007; Lau et al., 2008) and increasingly recognized to be central to syntax (Bornkessel-Schlesewsky & Schlesewsky, 2013; Matchin, Basilakos, et al., 2022; Matchin & Hickok, 2020; Pylkkänen, 2019), but this consisted of only a small number of voxels (5 total). Thus, it seems clear that these two violation types activated largely distinct resources in response to them. It is important to note here that, unlike ERP experimental designs, which time-lock the evoked response to the point at which a syntactic violation can be detected, the fMRI BOLD signal has dramatically less temporal resolution. This means that the differential networks activated by these two violation types may reflect processes occurring at different points in the timecourse of sentence processing, which cannot be effectively resolved in the present study.

The network observed for phrase structure violations corresponds well with the network that Meyer and Friederici 2016 identified as underlying the processing of grammatical sentences with non-canonical word orders (i.e., the subject and object appear in an order other than the default order for that language) and the processing of sentence containing embedded clauses through a meta-analysis of 20 articles. Meyer and Friederici link the activation of non-canonical word orders to a hypothetical process of re-ordering to retrieve the canonical word order (for semantic interpretation purpose), and they link the activation of embedded clauses to need to maintain independent mappings between the two subjects and the two objects in the sentence.

These processes very clearly involve working memory, whether domain-general or language-specific in nature (Fiebach et al., 2005; Matchin, 2018; Rogalsky et al., 2015); and indeed, this network has also been identified as a working memory network ((Buchsbaum & D’Esposito, 2008, 2019; Jonides et al., 2005; Majerus, 2019), see Buchsbaum et al., 2011 for a meta-analysis). Meyer and Friderici attempted to only include contrasts in their meta-analysis that controlled for non-syntactic working memory differences, such that this network could plausibly be interpreted as subserving syntactic (or syntactic/semantic) working memory processes. The meta-analysis identified two core regions: LIFG (BA 44) and left MTG/STG (BA 22).

The network observed for subject island violations corresponds well with the network that Binder et al., (2009) identified as the Semantic System through a meta-analysis of 120 articles that investigated semantic contrasts. To be included in that study, the contrasts had to be linguistic, and had to involve a difference in either the degree of access of stored semantic knowledge or the type of the semantic knowledge accessed. Crucially, contrasts were excluded if the semantic condition made greater demands on non-semantic systems, such as sensory, orthographic, phonological, syntactic, working memory, attentional, or response/motor systems. The contrasts tended to fall into three categories: words versus pseudowords, semantic vs phonological tasks, and high versus low meaningfulness (including meaningful vs nonsense sentences). We suspect that the focus on conceptual semantic contrasts likely means that parts of this network would be involved in most of the compositional semantic processes required by sentence processing, including building the thematic representation of the sentence. The meta-analysis identified 7 brain regions: 1) posterior inferior parietal lobe (AG and portions of SMG), 2) lateral temporal cortex (MTG and portions of ITG), 3) ventral temporal cortex (mid-fusiform and adjacent parahippocampal gyrus), 4) DMPFC, 5) IFG, 6) ventromedial prefrontal cortex (VMPFC), and 7) posterior cingulate gyrus. We note that this network also roughly corresponds to the semantic network theorized by Lau et al. (2008), including anterior temporal, inferior parietal, and anterior and posterior inferior frontal gyrus.

In order to evaluate the relationship between the activations obtained for PSV and subject island violations in our study and the systems involved in sentence complexity (i.e. clausal embedding)/noncanonicity (as identified by Meyer & Friederici, 2016) and the Semantic System (as identified by Binder et al., 2009), we obtained the activation maps associated with these previous meta-analyses (through personal communications with authors involved in these studies, Lars Meyer and Rutvik Desai, respectively) and computed the Dice coefficients between these maps and our uncorrected group level activation maps using AFNI. The Dice coefficient is a measure of spatial overlap between two or more thresholded brain maps (i.e., binarized statistical maps such that significant voxels are coded as 1 and non-significant voxels are coded as 0) which is used to assess the degree to which two maps are spatially similar to each other (Rombouts et al., 1999; Wilson et al., 2017). The Dice coefficient is calculated as 2 x V_intersection_ / (V_map1_ + V_map2_), where V_intersection_ reflects the number of overlapping voxels between the two maps, and V_map1_ and V_map2_ reflect the number of voxels in each of the two statistical brain maps being compared. The Dice coefficient ranges from 0 (no overlap) to 1 (perfect overlap). We found that there was a double dissociation in the Dice Coefficients: PSV vs. sentence complexity/noncanonicity was 0.20, subject island violations vs. sentence complexity/noncanonicity was 0.05, PSV vs. Semantic System was 0.04, and subject island violations vs. Semantic System was 0.14. Thus, while the the Dice coefficients were relatively low overall (which is not surprising for substantially different methodologies, different groups of participants, etc.), they patterned in accordance with our observations above.

Figure 6 plots the networks observed in the present study for PSV and subject islands in the top row, and the Binder et al. 2009 and Meyer and Friderici 2016 networks in the bottom row. In the following subsections, we discuss potential functional interpretations given the distinct networks activated by PSV and subject island violations.

**Figure 6.**
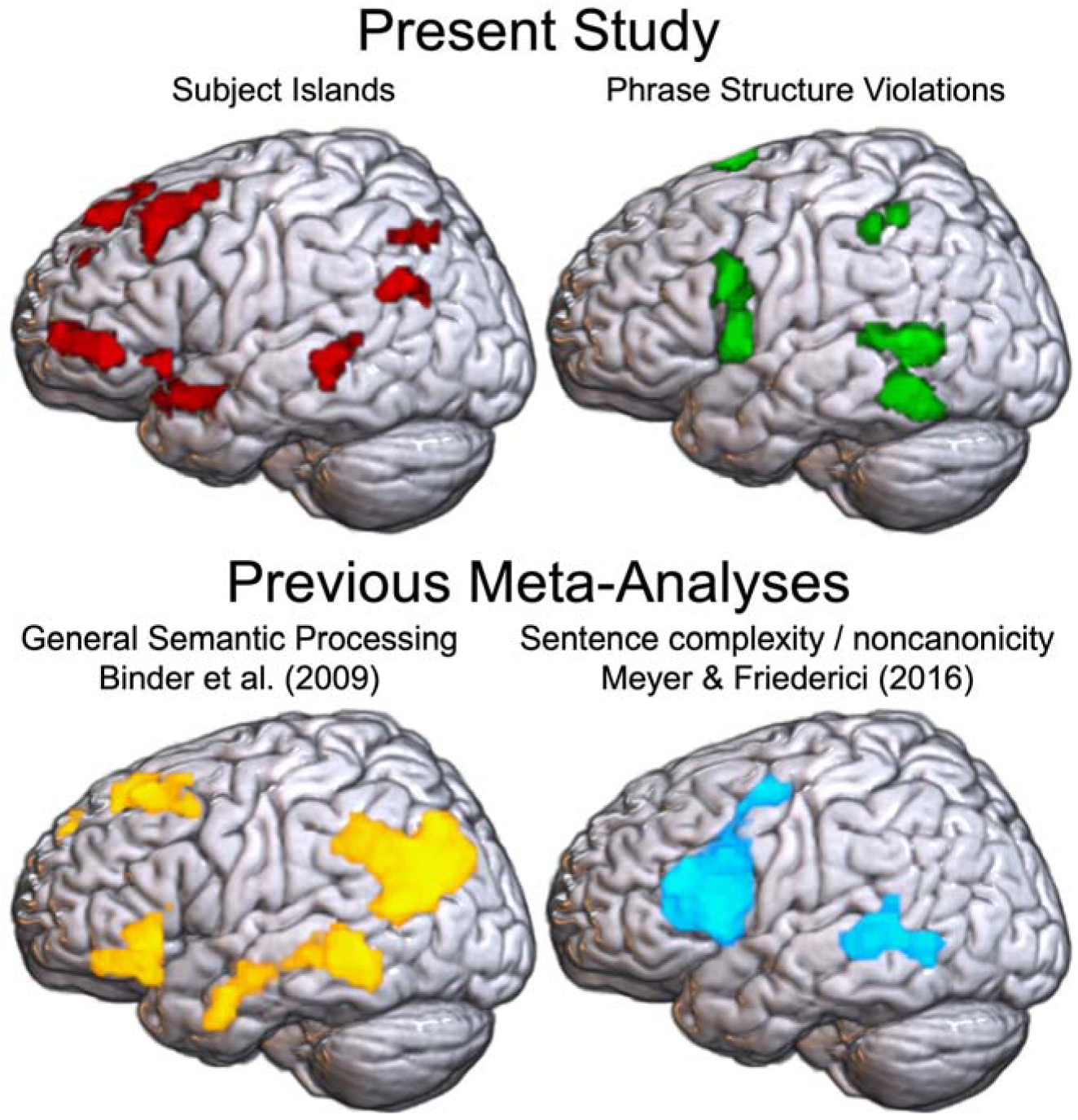
Comparison of the present results, main effect of subject island violations (TOP LEFT) and main effect of phrase structure violations (PSV) (TOP RIGHT) with previously published meta-analyses, general semantic contrasts (BOTTOM LEFT) (e.g. real words > pseudowords, semantic decision task vs. phonological decision task) (Binder et al., 2009), and sentence processing contrasts based on presence of clausal embedding or noncanonical word order relative to canonical word order (BOTTOM RIGHT) (e.g. object-relatives > subject-relatives) (Meyer & Friederici, 2016). Data from Binder et al. (2009) and Meyer & Friederici (2016) reproduced with novel visualization, with thresholded results binarized.

### 5.2 A functional interpretation of the activation pattern for phrase structure violations

Recall from section 2.2 that there is general consensus in the literature that violation- identification processes for PSV are likely related to local (syntactic) predictive mechanisms and that the repair processes are likely related to reordering and working memory – i.e., a search for a grammatically licit order of the items in memory. Given that our results appear to show activation of a reordering/WM network, it seems possible that our results are driven by the repair processes. We also note that PSV did not appear to activate the semantic network from Binder et al. 2009. This suggests that the repair processes did not create a heavier semantic burden on the violation condition relative to the control. One plausible interpretation of this is that the end result of the repair process is a sentence with the same meaning as the control condition.

Though the goal of our study was not to identify the source of ERP results, we do think it is valuable to explore how our results could be linked back to the Neville et al., 1991 ERP results. Recall that Neville et al. observed both a LAN and a P600 for PSV. They interpreted the LAN as indexing the violation identification processes and the P600 as indexing repair processes. If we adopt the same analysis, then the activation that we observe would be most likely related to the P600. This would suggest that at least some P600s might have a source in the reordering/WM network. However, we could also adopt a different analysis in which the LAN does not index violation detection, but rather both the LAN and P600 index repair processes. Under that analysis, it is possible that the LAN could index the working memory component of the reordering process, while the P600 indexes a later stage, such as the identification of a grammatical sequence. In that case, the activation of the reordering/WM network that we observed would be most likely related to the LAN. Though we can’t adjudicate between these two analyses based on our fMRI results alone, the existence of these two relatively constrained possibilities suggests that there could be theoretical value to future studies investigating the sources of the LAN and P600 in phrase structure violations.

Before concluding this subsection, we should also note that our results may appear to conflict with the results of a recent fMRI study, Mollica et al., 2020, which found that word order permutations similar to our PSV condition did not produce increased activation compared to conditions without such permutations, specifically in language-related brain regions. In that study, activation for word order permutations only became distinguishable for sentences that contained a large number of permutations, resulting in a decrease of activity relative to fully grammatical sentences. Crucially, Mollica et al. first localized language-related regions by comparing sentences to unstructured lists of pseudowords and then used these regions as ROIs. These ROIs overlap with the networks that we observe here, but may diverge at least to some degree, so the lack of alignment between our results could be partially an artifact of the choice of ROIs. Potential evidence for this lies in the fact that Mollica et al. did report that regions adjacent to their ROIs showed elevated activation to increased word order permutations, including a cluster straddling the posterior IFG/precentral gyrus that seems similar to parts of the network identified in our uncorrected whole-brain analysis of PSV. Additionally, another crucial factor is likely to be the fact that in our study, 50% of sentences did not have a word order permutation, and 50% had a simple linear word order permutation, whereas in Mollica et al., only 20% of sentences in the critical experiment were grammatically intact, 20% of sentences contained a simple word order permutation as in our study, and 60% of sentences had increasingly more complex permutations. Given the large body of previous studies that have identified robust activations similar to what we observed here for phrase structure violations based on word order permutation (Embick et al., 2000; Friederici et al., 2003; Kuperberg et al., 2000, Meyer et al., 2000; Moro et al., 2001; Newman et al., 2001; Rüschemeyer et al., 2005), we suggest that differences in methodology largely account for these seemingly discrepant findings.

### 5.3 A functional interpretation of the activation pattern of subject island violations

Recall from section 2.2 that the three classes of island theories might be plausibly interpreted as making distinct predictions about the identification and repair processes triggered by island violations. For syntactic theories, the identification processes would likely be related to long-distance dependency processing and gap identification, perhaps implicating working memory processes. The repair processes would likely be related to the thematic structure of the sentence, such as incomplete argument structure at the gap location and vacuous quantification associated with the filler. For information-structure theories, both the identification and repair strategies would be related to a focus/backgroundedness clash. Finally, for processing theories, both the identification and repair processes would be related to working memory overload. Crucially, we observed activation in areas that correspond roughly with the semantic system network identified by Binder et al. 2009 (see also the meta-analyses in Hagoort & Indefrey, 2014 and Hodgson et al., 2021), with almost no overlap with the network activated by PSV, which appeared to be a WM/sentence reordering system. This pattern appears to be consistent with the repair processes (but not identification processes) that are plausibly predicted by the syntactic class of island theories. This pattern also appears to be inconsistent with the predictions of processing-based theories of island effects. But one lingering question is whether it is compatible with the identification and repair processes that are plausibly predicted by the information-structure theories.

There is substantial overlap between the semantic system and the “default network” that is typically identified using resting-state (and task-based) functional correlation data (Binder, 2012; Buckner & DiNicola, 2019), as well as brain networks for theory-of-mind processing (Saxe, 2006; Saxe & Kanwisher, 2003; Shain et al., 2023; Whitfield-Gabrieli et al., 2011). The significant overlap between the semantic system, the default mode network, and the theory of mind network raises the possibility that semantic processing, internal planning, ascertaining communicative intent, and pragmatic reasoning may be in part supported by a shared brain system. In our search of the literature, we could identify only one previous fMRI study that specifically investigated focus/background clashes - van Leeuwen et al., (2014), which found effects that partially overlapped with these systems. While we have interpreted our subject island results primarily in terms of the semantic system and reanalysis of the conceptual-semantic interpretation of the sentence, it is possible that information-theoretic accounts might predict the present results if the activations reflect increased reliance on assessing belief and communicative intent.

It is, of course, also possible that the apparent activation of the semantic system network is indicative of a novel semantic analysis of either the identification or repair processes triggered by subject island violations. There are semantic theories of island effects (e.g., Abrusán, 2011; Szabolcsi & Zwarts, 1993) that seek to explain island effects as an incompatibility between the semantics of a wh-dependency and the semantics of the island structure. However, these have never, to our knowledge, been extended to subject islands. This is because the current versions of semantic theories rely on the existence of a semantic operator within the island (such as a question operator for wh-islands or a negative operator for negation islands). It is unlikely that there is any such operator for subjects. This is why we did not consider semantic theories in the review in section 2. We only note them here to say that if one were inclined to interpret the activation of the semantic system network as evidence that the source of subject island violations is semantic, it would require postulating a novel semantic theory of subject islands. We leave that as a possibility for future research.

As with PSV, we do think it is valuable to explore how our results for subject island violations could be linked back to the Neville et al. 1991 ERP results. Recall that Neville et al. observed an increase in P2 amplitude (likely related to predictive processing) and a P600 for subject island violations. It seems more likely that the activation pattern that we observed in fMRI is related to the P600 effect than the P2 effect. Neville et al. interpreted the P600 as reflecting repair processes, and that is in line with the conclusions that we drew from the interaction of the semantic nature of the network and the possible predictions of the three existing classes of theories of island effects. If our logic is on the right track, then the semantic system network is a possible source for the P600 to subject island violations.

### 5.4 Potentially distinct neural sources for the P600 responses to phrase structure and subject island violations

Given our fMRI results, we have postulated distinct neural sources for different P600 effects, in line with previous suggestions (Hagoort, 2005; Neville et al., 1991). Our study was not designed to evaluate ERPs directly, so here we simply note that the distinct patterns of results that we observed for phrase structure and subject island violations suggest that there may also be theoretical value in exploring the neural generators of P600 effects to distinct syntactic violations. More functional neuroimaging research is needed to confirm these speculations.

Alternatively, the apparent contrast between our results and those of Neville et al. (1991) could be due to differences in stimulus presentation. In the Neville et al. (1991) study, 50% of their stimuli were acceptable, each violation types only constituted 1/8 of the total stimuli, a ratio of 7 to 1. By contrast, violations were highly frequent in our study: only 25% of the sentences presented were acceptable, and both PSV and subject islands were presented in 50% of the stimuli (due to the fact that 25% of stimuli were both PSV and subject island). The P600 has been shown to disapper when violations are extremely frequent (80% in Hahne & Friederici, 1999), or when participants are not consciously aware of the violation (Batterink & Neville, 2013). Thus, it is unclear the extent to which an ERP study based on our design would in fact elicit a P600 effect. In that case, our fMRI results might not be tracking the P600 at all. A high rate of violation sentences may be an advantage in fMRI studies seeking to use brain data in order to differentiate the sources of different violation types. Repeated presentation may be an effective method for using functional neuroimaging to discover the underlying issue with grammatical violations by limiting potential confounding generalized oddball effects. But, differences in the ratio of violations may also complicate the comparison of fMRI and ERP responses to vioations.

A related issue concerns the unusually high acceptability of Subject Island violations in our study compared to previous studies (Sprouse et al., 2011, 2012, 2016), which we believe is related to the proportion of sentences containing violations. Because the higher acceptability of Subject Islands makes them in some sense more similar to the Grammatical condition than PSV, it is logically possible that an fMRI study which obtained lower acceptability for Subject Islands would show greater overlap with PSV or other syntactic violations. However, the fact that Subject Islands produced substantial activation relative to the Grammatical condition suggests that participants did not view them as identical to each other; and the fact that this activation pattern is distinct from the activation patterns for PSV in turn suggests that Subject Islands involve some qualitatively distinct processes from PSV.

### 5.5 Limitations and future directions

One major limitation of our study is the relatively low statistical power of the results we obtained: while the main effect of subject island and the main effect of PSV produced distinct significant activations, we also relied on the uncorrected statistical maps for inference regarding the broader networks activated by these conditions. One possible explanation for the low power is that the violation effects themselves could be relatively small. We note that subject island violations have never, to our knowledge, been reported using fMRI; and even within the ERP literature, we know of no replications of the Neville et al. 1991 results, so it is difficult to assess the expected size of the effect. Another possibility is that a relatively high proportion of sentences contained a violation of some kind (75%). Though this is a necessary consequence of testing violation paradigm along with an articulatory rehearsal paradigm, it could have led to a saturation effect, both because participants expected violations to occur, and because of the well-known observation that the BOLD response to a particular stimulus characteristic decreases with repetition (Grill-Spector et al., 2006; however, this is mitigated by the fact that the two violations did not appear to have the same source). Another possibility is that participants may have used a more formulaic processing strategy than in natural settings because every sentence contained a similar filler-gap dependency with a second clause introduced by the verb *think*. Future studies could address these issues by increasing the structural variability among experimental materials, increasing the number of non-violation sentences (perhaps by sacrificing other comparison tasks), or even by augmenting a natural story to include the violations of interest.

In addition, our individual participant analyses showed substantial variability in terms of the activation patterns for these two violation types, which likely contributed to the limited statistical robustness (Thirion et al., 2007). However, these individual participant analyses also revealed that, with one exception, there were largely distinct activation patterns for PSV and Subject Islands. Given that group-level analyses can obscure the true underyling spatial distribution of effects in individual participants (Burton et al., 2001; Fedorenko & Kanwisher, 2009), particularly in association cortex (Frost & Goebel, 2012; Tahmasebi et al., 2012; Vázquez-Rodríguez et al., 2019), these individual participant analyses help to bolster our conclusions about the separability of the activation patterns elicited by these violation types, and suggest that future research in this area incorporate both functional localizers and individual participant analyses.

Another issue concerns the use of reverse inference: we have interpreted the nature of the linguistic violations in light of apparent activation in brain networks that have been previously characterized in functional neuroimaging research. However, there are important issues with reverse inference, namely that the validity of these inferences depends on both the confidence of the cognitive processes supported by those brain networks, as well as the confidence that the activations we obtained here indeed align with those same brain networks (Poldrack, 2006). The present study is certainly subject to strong limitations on both of these points. Future work should incorporate functional localizers to help strengthen the confidence of conclusions that can be drawn.

Despite the limitations of this study, we believe that our experimental approach and tentative results could be followed up with new research directions. There is potential theoretical value in testing multiple distinct violation types within the same experiment, and using the functional interpretations of the activation networks from the cognitive neuroscience literature to adjudicate among the potential identification and repair processes derived from the linguistics and psycholinguistics literature. More specifically, comparing PSV systematically to other violation types that have no obvious repair by reordering (like subject islands) could help determine whether the network revealed in the meta-analysis by Meyer & Friederici is specific to reordering. Similarly, it could also be revealing to systematically test different island violation types, to see whether they all activate the Binder et al. semantic system (as potentially predicted by the syntactic approaches to islands) or whether they might trigger distinct repair processes (and therefore might have distinct sources). Finally, at the narrowest level, it might be revealing to explore specific subcomponents of the semantic repair processes that could underlie subject island violations, perhaps by investigating vacuous quantification in isolation, or by investigating missing thematic arguments in isolation. This could reveal subnetworks within the semantic system. Similarly, it might be valuable to explore a wider array of information-structure manipulations to try to identify the network that is specific to focus/backgroundedness calculations (and/or clashes). However, such approaches must carefully incorporate several methodological considerations, such as the number of violation trials relative to fully acceptable sentences, sufficient statistical power, and functional localizers to independently verify the relevant sentence processing/reordering and conceptual-semantic networks.

## 6. Conclusions

Our experiment illustrates that neuroimaging data can provide complementary insight into questions about syntactic theory that are difficult to address using traditional methods (i.e., acceptability judgments). There are two key observations a that can be made from the results of this study. The first is that there was little common activation between island violations and PSV, with the little overlap occurring in posterior temporal cortex that is unlikely due to executive function resources. This suggests that there was no common generic “violation” response for these two conditions. Future studies could harness these or other violations in order to more closely match neuroimaging data with the bread and butter of syntactic theory, which are minimal pairs of acceptable and unacceptable sentences. Critically, such studies should be certain to include multiple violation types. The second is that PSV and subject island violations activated distinct brain networks with established functional interpretations: PSV activated brain regions previously implicated in reordering working memory (also supported by the overlap of PSV and articulatory rehearsal), and subject island violations activated brain regions previously implicated in conceptual-semantic processing. These functional interpretations allowed us to draw tentative conclusions about the processes triggered by these violations, how the processes map to existing ERP results, and about the grammatical source of the violations. Though each of these conclusions needs to be confirmed with dedicated studies, we believe this approach could help form links across linguistics, psycholinguistics, and neurolinguistics.

## 7. Data Availability Statement

Data will be made available upon reasonable request through contact with the lead author.

## 8. Author Contribution

William Matchin: Formal Analysis; Writing – Original Draft; Writing – Review & Editing; Visualization

Diogo Almeida: Conceptualization; Investigation; Data Curation

Gregory Hickok: Resources; Supervision; Project administration; Funding acquisition Jon Sprouse: Conceptualization; Investigation; Methodology; Formal Analysis; Writing –

Original Draft; Writing – Review & Editing; Visualization; Supervision; Project Administration

## Acknowledgements

We would like to thank Rutvik Desai for original data from Binder et al. (2009) and Lars Meyer for original data from Meyer & Friederici (2016). We would also like to thank the audiences of the California Meeting for Psycholinguistics (CAMP) 2017 and the CUNY sentence processing conference (now Human Sentence Processing, HSP) 2018 for their feedback on this work.

Finally, we would like to express deep gratitude to Cory Shain for comments and feedback on a previous version of this manuscript.

## 9. Funding Information

This work was supported by NIH NIDCD grant DC03681 awarded to Gregory Hickok.

## Notes

### Competing Interest Statement

The authors have declared no competing interest.

### Summary of Updates

new individual subject analyses have been added, introduction and discussion have been updated to modulate the claims and update the literature review, additional details about stimulus materials have been added

